# 8-channel Tx dipole and 20-channel Rx loop coil array for MRI of the cervical spinal cord at 7 Tesla

**DOI:** 10.1101/2023.02.08.527664

**Authors:** Nibardo Lopez Rios, Kyle M. Gilbert, Daniel Papp, Gaspard Cereza, Alexandru Foias, D. Rangaprakash, Markus W. May, Bastien Guerin, Lawrence L. Wald, Boris Keil, Jason P. Stockmann, Robert L. Barry, Julien Cohen-Adad

**Author notes:** Correspondence Nibardo Lopez Rios. NeuroPoly Lab, Institute of Biomedical Engineering, Polytechnique Montréal, 2900 Edouard-Montpetit Bld, H3T1J4, Montreal, QC, Canada.

## Abstract

The quality of cervical spinal cord images can be improved by the use of tailored radiofrequency coil solutions for ultra-high field imaging; however, very few commercial and research 7 Tesla radiofrequency coils currently exist for the spinal cord, and in particular those with parallel transmit capabilities. This work presents the design, testing and validation of a pTx/Rx coil for the human neck and cervical/upper-thoracic spinal cord. The pTx portion is composed of 8 dipoles to ensure high homogeneity over this large region of the spinal cord. The Rx portion is made of 20 semi-adaptable overlapping loops to produce high Signal-to-noise ratio (SNR) across the patient population. The coil housing is designed to facilitate patient positioning and comfort, while being tight fitting to ensure high sensitivity. We demonstrate RF shimming capabilities to optimize B_1_^+^ uniformity, power efficiency and/or specific absorption rate (SAR) efficiency. B_1_^+^ homogeneity, SNR and g-factor was evaluated in adult volunteers and demonstrated excellent performance from the occipital lobe down to the T4-T5 level. We compared the proposed coil with two state-of-the-art head and head/neck coils, confirming its superiority in the cervical and upper-thoracic regions of the spinal cord. This coil solution therefore provides a convincing platform for producing the high image quality necessary for clinical and research scanning of the upper spinal cord.

## Introduction

Clinical and research MRI studies of the cervical spinal cord can benefit significantly from the increased signal-to-noise ratio and contrasts resulting from the recent development in ultra-high fields. The high resolution achieved, particularly at 7 Tesla (T), has great potential for enhanced structural, functional, quantitative and spectroscopic studies of the small structures that comprise this region^1,2^. The anatomical characteristics of the spinal cord present a challenge for magnetic resonance imaging (MRI) radiofrequency (RF) coils to produce both a uniform transmit field and a high receive sensitivity at 7 T. The confluence of the large inferior-superior dimension of the spinal cord, its dissimilar curvature and posterior location, and the variance in surrounding body circumference along its longitudinal axis, has inspired the development of a wide range of tailored solutions for different sections of the spine.

Overall, spine coil transceiver archetypes can be classified based upon the anatomical region to which they are tailored: head-neck, cervical spine, thoracic, and lumbar. Head-neck coils^3–5^, as well as cervical spine coils^2,6^, are typically conformed to the complex shape of these sections of anatomy—the focal coverage produces higher transmit efficiency and receive sensitivity. Imaging of the thoracic and lumbar regions requires enlarged coverage in the superior-inferior direction, which has been accomplished with planar arrays residing posterior to the subject^7–11^. Torso coils are also used for these two regions, being either rigid^12^ or made flexible^13–15^ to contend with the variance in body size between subjects. In a select number of institutions, whole-body coils have been developed^16,17^ that reside outside the bore liner—this engineering feat would allow for imaging of the entire spine.

There exists a dearth in the literature for coils tailored to an imaging region spanning from the occipital lobe to the mid-thoracic—a region probed for the detection of multiple sclerosis lesions^18^. The disparate requirements for conforming the receive coil to the vastly different size and shape of the neck versus the upper torso must be accommodated to optimize sensitivity: a tailoring of the receive-coil geometry to this anatomy is therefore imperative. Conversely, a transmit coil must be sufficiently distant from the body to produce a uniform field, yet not so distant as to produce inadequate efficiency for attaining the required RF pulses (a common challenge for 7 T spine coils)—this all must be achieved while avoiding the geometric impediments of the shoulders and the receive coil.

At 7 T, spine coils have accomplished these tasks with either transceive arrays^4,10,11^ or separating transmit and receive functions into disparate coils^2,3,5–7,16,19,20^. This latter approach often permits a greater number of receive elements to be employed, and as such was chosen as the topology for the current study.

Receive arrays must be both highly sensitive and have the requisite geometrical layout to reduce noise amplification during the reconstruction of accelerated images^21,22^. Various challenges must be overcome during the design of a receive coil for the target region of interest, some of which are specific to high-field imaging. Among these confounds is the location of the spinal cord, where the signal-to-noise ratio (SNR) exhibited by existing arrays is normally reduced^23,24^. The spinal cord’s depth varies along its length, a challenge that is exacerbated by the variability of the human anatomy when designing a coil to accommodate a large portion of the population. Consequently, to maximize signal sensitivity, it is advantageous to minimize the distance between receive elements and the spine. This results in coil elements being distributed over a complex surface, with a high density of elements proximal to the neck (thereby increasing coupling between elements) and a low density in the upper thorax, including no coverage around the shoulders (causing a signal reduction). In this regard, adjustable designs^25^ can be beneficial, despite their increase in design complexity. Likewise, the array can be physically split to facilitate accommodation of the subject and to reduce stress during scanning. The resultant complexity in the coil former leads to a more difficult decoupling process due to the diversity of element dimensions and layouts. Furthermore, the proximity and geometry of the transmit coil must be considered to determine the location of the preamplifiers and receive-coil cabling that limits transmit-receive interaction.

Transmit coils are responsible for creating transverse magnetic fields (B_1_^+^) that are uniform, power efficient and specific absorption rate (SAR) efficient—challenges exacerbated at high field strengths due to the reduced wavelength within the human body and the resultant complex wave behavior^26,27^. Improvements in B_1_^+^ uniformity^28,29^, required to mitigate spatially dependent tissue contrast, have been addressed through the advent of parallel-transmit coils^30,31^ augmented with RF shimming^32,33^ and spatially selective RF pulses^34^. Power efficiency must only be sufficient to produce the requisite RF pulses within the constraints imposed by the RF power amplifiers; therefore, the more salient performance metric at high field is SAR efficiency, as local SAR is often the practical limitation when creating pulse-sequence protocols.

For spinal cord imaging at ultra-high field, various design topologies have been developed for transmit coils. Achieving sufficient transmit efficiency in the spinal cord—a relatively deep structure—can prove challenging. Local transmit coils can reduce this difficulty by placing the coil in close proximity to the subject. This has been accomplished with two-channel coils^7,35–37^ and array coils^2,5,8,20,38–40^. To additionally leverage the capability of array coils to improve transmit uniformity through RF shimming or spatially selective pulses, an 8-channel transverse electromagnetic (TEM) coil^17^ and a 16-channel meandered stripline coil^41^ were developed for torso imaging, including the lower spinal cord. A more general solution for whole-body imaging has been to build the transmit coil into the bore liner: these have included a TEM coil^17^ and 32-channel stripline array^42^. The latter solution combined the capacity of a large-diameter, high-density transmit coil (to improve uniformity) with a custom power amplifier for each transmitter near the magnet, thereby compensating for the commensurate reduction in efficiency.

Recently, the dipole has been introduced as a constituent element for ultra-high-field transmit coils^43–46^ due to being an efficient high-frequency radiator—thereby improving efficiency in deep tissues—while commensurately reducing SAR^47,48^. As such, dipoles have been used as the building block for transmit coils intended for head imaging^49–52^ and body imaging^15,53^. For spine imaging, dipoles have been implemented to improve transmit efficiency^7,12,39^, SAR efficiency^19,54^, and to increase the potential field-of-view^36^.

In this work, we present a single-housing transmit/receive coil, composed of eight geometrically adapted Tx dipoles and 20 anatomically shaped Rx loops, for an extended field-of-view spanning from the occipital lobe to the upper-thoracic spine (T4-T5). The coil features an adjustable anterior section that facilitates the positioning of subjects and allows the Rx array to be adapted to them. The coil was validated by simulations, on the workbench and in the scanner with multiple subjects of variable morphologies. RF shimming was demonstrated with the multi-channel Tx design to further improve the *B*_1_^+^ uniformity, power efficiency, or SAR efficiency. The coil was compared to a commercially available coil and a recently developed head-neck coil^5^.

## Methods

### Coil construction and evaluation on the bench

The Tx and the Rx coils were mounted on a single structure with a fixed posterior section and a hinged anterior section to expedite the setup and removal of subjects (**Figure 1A,B**). Fifteen receive elements were mounted on the close-fitting, 3-mm-thick inner surface of the posterior section to produce high signal sensitivity. In this fixed section, transmit elements Tx-1 to 7 were installed on machined polycarbonate sheets: three posterior, and two each on the left and right sides. Tx-8 and Rx-16 to 20 were placed on the anterior section. These five receive elements compris a functionally independent assembly whose distance to the subject can be manually adjusted by the MR technologist, or ultimately detached, allowing the coil to be adapted to an extended population. The anterior section is locked in place during scanning to prevent movement. External cabling enters the posterior section of the coil and cannot be in direct contact with the subject during scans. The cable of the Tx-8 element was externally routed from the fixed to the hinged section and protected with cable wrap. A mirror, with adjustable Z-axis position and tilt angle, was integrated underneath the anterior transmit housing to reduce patient anxiety and to allow for the presentation of visual stimuli. Open spaces around the face of the subject were maximized to further reduce claustrophobia. A head-neck phantom (**Figure 1C**) was also built to adjust and evaluate the coil.

**Figure 1.**
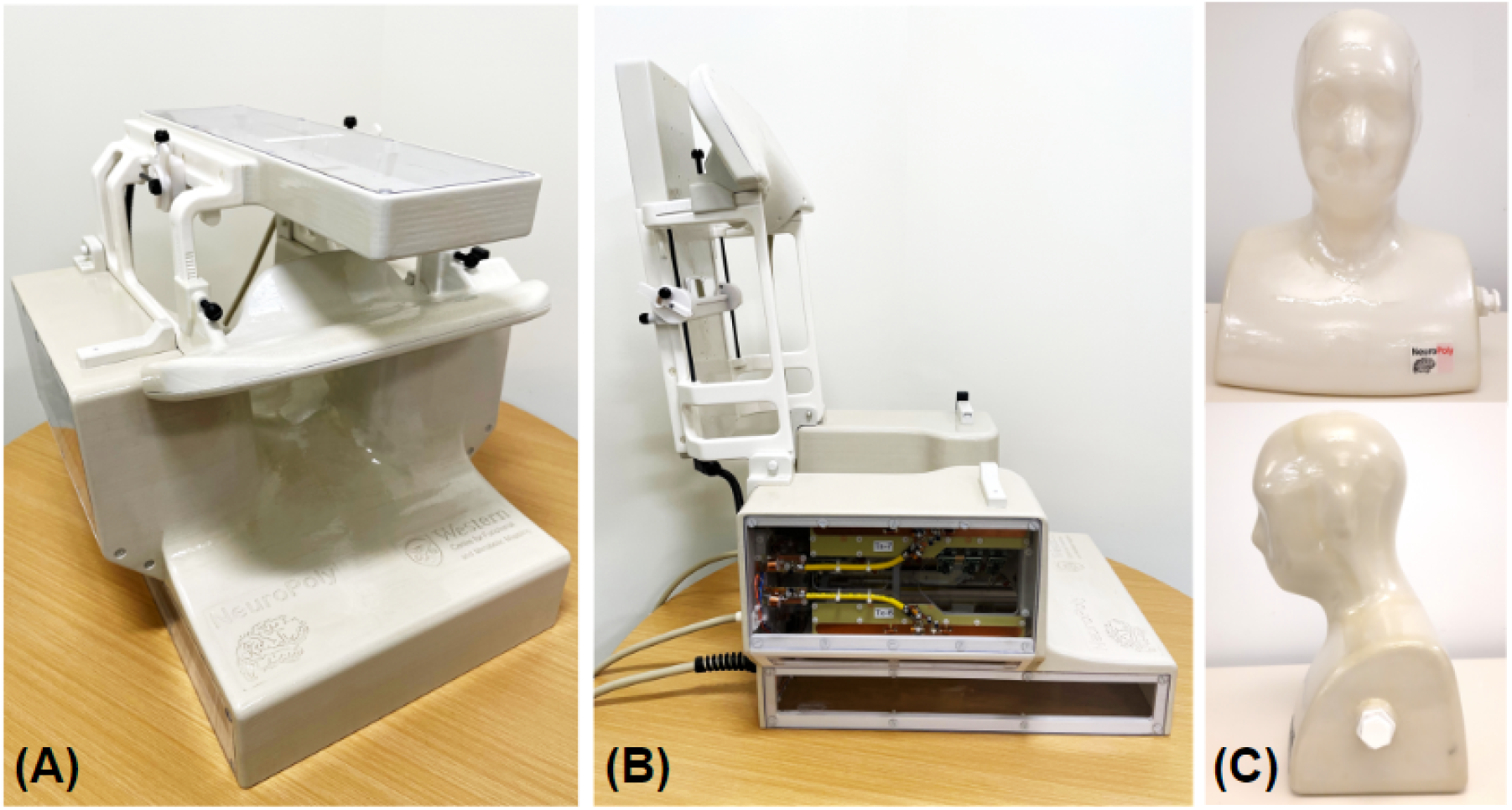
Views of the assembled Tx/Rx coil in its imaging mode (**A**) and with the hinged assembly (Tx-8/Rx-16-20) completely open to allow subject placement (**B**). The head-neck phantom built for coil adjustment and evaluation is shown in (**C**).

#### Receive coil

We used a computer-aided design tool (Autodesk, AutoCAD, Version 2018, San Francisco, CA) to design a close-fitting inner coil former. As a reference for this process we utilized a 3D average head model created by a procedure described in an earlier work^55^. The basic structure of the resulting former was a complex surface that slightly reduced the typical curvature of the cervical spine during scans, while leaving additional space for padding. The anterior of this surface was modified from the average head model to facilitate its adaptability to the highly variable anatomy of that region over the population. Based on the 3D probabilistic localization of the spinal cord obtained with the procedure cited above, we determined that 20 circular loops placed directly on this surface could image the entire region of interest with an adequate sensitivity (**Figure 2A, C**). The loops located in the fixed section (Rx-1 to 15) were proximal to the posterior and lateral portions of the neck, while extending from the occipital to the mid-thoracic regions. The remaining loops (Rx-16 to 20) were intended to improve the overall sensitivity of the coil on the anterior portion of the neck and the upper chest, especially in the inferior aspect of the region of interest, where the distribution of the posterior 15 receive elements was less dense.

**Figure 2.**
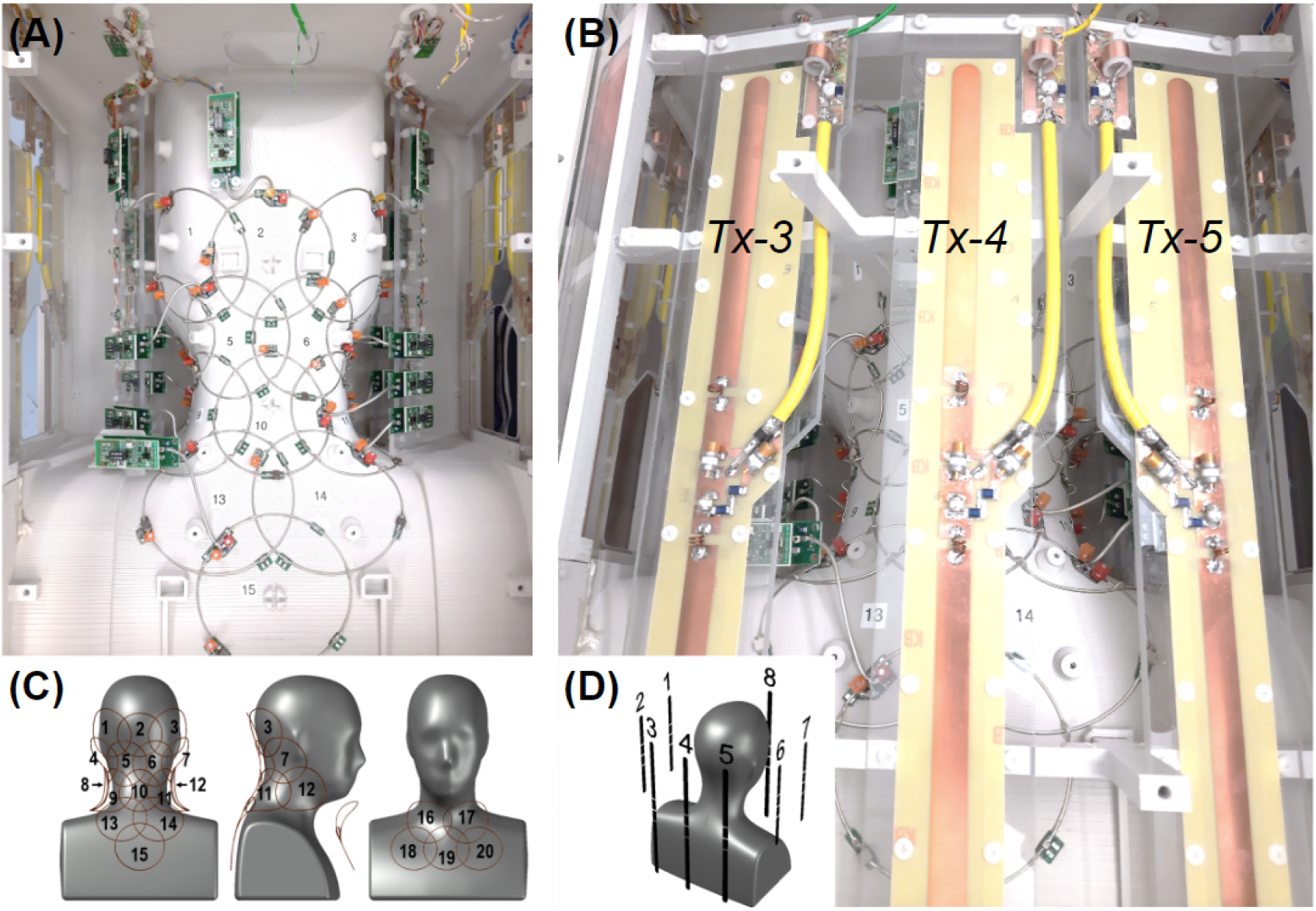
Posterior views of the coil and element layout. (A) Rx subarray comprising elements 1 to 15; lateral Tx elements can be seen on the sides: 1, 2 on the left and 6, 7 on the right. (B) Full view of Tx elements 3, 4 and 5. (C) Distribution of Rx loops on the posterior (1-15) and anterior (16-20) Rx coil sections. (D) Posterior-right perspective view of the phantom showing the arrangement of Tx dipoles.

The receive-loop contours were generated in CST Studio Suite (Waltham, MA, USA) using the previously designed coil sections imported from AutoCAD. To do this, spheres were sequentially created and intersected with the former sections. The lines defining the intersections between the formers and the spheres were converted into wires (r = 0.64 mm) resulting in circular loops. Each loop was segmented into four sections of approximately equal length, creating four gaps where simulation ports were connected. Tuning and matching of the loops were performed by co-simulation after assigning variable components to the simulation ports. Coupling between adjacent pairs was minimized by adjusting the positions and diameters of the corresponding spheres. Final sphere diameters were between 88 and 98 mm. The symmetry of the coil was leveraged to significantly reduce the number of iterations. Finally, the sensitivity profile of the complete array was estimated by computing and combining individual B_1_^-^ maps.

The final array structure was exported back to AutoCAD and subtracted from the original coil formers. This operation generated grooves that were used as a guide to lay the loops. Loops were made of 16 AWG tinned copper wire (Belden, 8013) and were incorporated into an electrical design with active and passive detuning mounted over different tuning capacitors (**Figure 3A**). A variable tuning capacitor was used for initial adjustments then replaced with a fixed component to increase coil reliability. To increase coil safety, all loops were provided with fast non-magnetic fuses (Siemens Healthineers, Erlangen, Germany) with a rating of 315 mA at the operating frequency. A 65-mm-long semi-rigid coaxial cable (Carlisle, UT-85C_FORM) allowed for separating preamplifiers from loops to minimize the noise figure, increase stability, and reduce the interaction with Tx elements. Cable traps were integrated into the output cables to reduce undesirable crosstalk between receive channels and coupling with the Tx-coil. The selected preamplifiers (Siemens, 10185 702 E2) also had internal cable traps between the first and second amplification stage.

**Figure 3.**
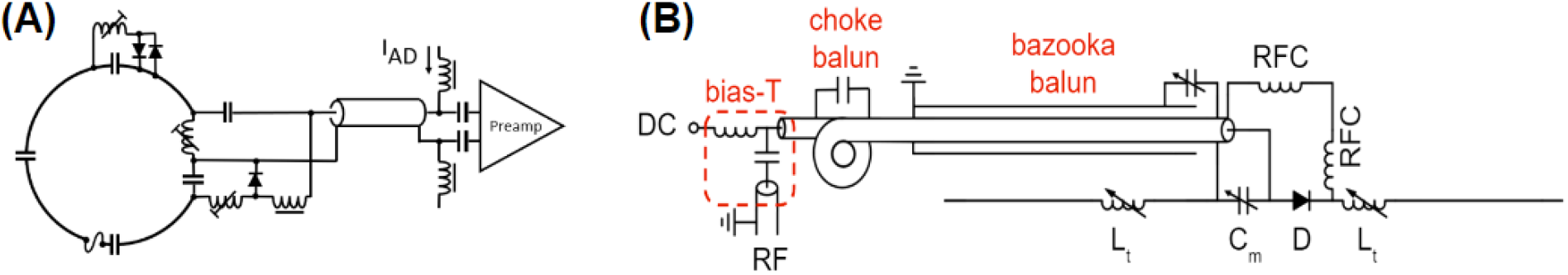
Electrical schematic of the Rx (**A**) and Tx (**B**) elements.

After assembling the receive coil and adjusting all active and passive detuning circuits, the output impedance of the Rx loops was set as close as possible to 75 Ω to optimize the noise match of the selected preamplifiers. Coupling was subsequently evaluated by measuring the transmission coefficient between all possible loop combinations for both the posterior and the anterior subarrays. For this purpose, 75-to-50 Ω transformers were inserted between the input connectors of the preamplifier socket boards and the VNA probes. The Tx coil was not installed during these tests to facilitate access to the socket boards, as it was found to cause negligible effects on the Rx coil parameters.

#### Transmit coil

The transmit design was based on full-wave electromagnetic simulations performed in CST Studio Suite. Eight dipoles of two different lengths were selected as transmit elements. Each dipole was modeled as having three ports: two ports representing tuning inductors and one central port representing a parallel matching capacitor. Tuning and matching was accomplished by means of a co-circuit simulation.

The B_1_^+^ distribution within the phantom was simulated to compare to experimental results and to subsequently validate the coil model. Simulated B_1_^+^ maps of individual transmit elements were scaled to equate the mean phase and 95^th^ percentile of their magnitude to experimental B_1_^+^ maps, as described in an earlier work^56^, ensuring agreement between simulated and experimental maps. The simulated and experimental B_1_^+^ maps of individual transmitters were combined in circularly polarized (CP) mode and with 10,000 different random RF shim settings: the difference between the predicted combined B_1_^+^ maps (mean and 95^th^ percentile) was calculated to quantify their agreement over a large range of shim solutions.

With the coil model validated through phantom simulations, four CST body models were simulated (Hugo, Gustav, Laura and Donna) for use in online SAR monitoring. Local SAR matrices (Q-matrices) were derived for each body model and catenated prior to compression into virtual observation points (VOPs)^57^: the catenation of multiple body models of varying size and position ensured that the worst-case scenario for local SAR would be implemented on the scanner. To account for additional inaccuracies in coil modeling, differences in the population and inaccuracies in the online measurement^58^, a conservative safety factor of 2.0 was added to the SAR prediction.

To construct the transmit coil, the dipoles were machined from copper-clad Garolite. All dipoles were 1.6-cm wide with rounded ends to mitigate high conservative electric fields. **Figure 2B** shows the structure of the posterior Tx elements. Anterior and posterior elements were 38.1-cm long to provide adequate coverage from the occipital pole to the fourth thoracic vertebrae. Alternatively, the length of the lateral dipoles was set at 22.1 cm, in practice to avoid the impediment of the subject’s shoulders; however, a commensurate benefit to offsetting their B_1_^+^ profiles with respect to the longer dipoles was an enhanced ability to RF shim over the longitudinally oriented spinal cord. The distance between dipoles (a minimum of 10 cm) was chosen to permit a level of coupling that could be compensated for with RF shimming. The distance between any dipole and the center of the imaging region (C3) ranged from approximately 14 to 22 cm—this distance was a compromise between transmit uniformity and efficiency. The sparse geometry of transmit dipoles (see **Figure 2D**) was also selected to allow receive-coil preamplifiers to be placed in regions that would reduce interaction between the constituent coils. Thus, the lateral transmit elements were placed 5 cm from the nearest preamplifier to mitigate coupling and the corresponding adverse effects to coil performance.

Two high-Q inductors (approximately 10 - 80 nH, depending on dipole length and load) were incorporated into each dipole to electrically shorten them to resonate at 297.2 MHz. The electrical design is presented in **Figure 3B**. Dielectric coupling between the ends of the posterior (long) dipoles and the supports caused their resonant frequency to shift by approximately 55 MHz; therefore, the housing beneath Tx-8 and the polycarbonate supports beneath Tx-3 to 5 were hollowed out to decrease the frequency shift and allow for these elements to be tuned to the correct frequency. Although this was also done for the shorter dipoles for mechanical reasons, it was not necessary as they required a larger tuning inductance.

An electrically shortened bazooka balun (Belden 9222 triaxial cable, Johanson 59H01 high-power variable capacitor) and a hand-made cable trap (distal to the bazooka balun and dipole input) were installed between each dipole and its transmission line to attenuate common-mode RF currents. A parallel matching capacitor (Johanson 59H01, 1-25 pF, rated to 1000 V_DC_) was employed to match dipoles to 50 Ω. Active-detuning was achieved with a serial high-power PIN diode (Chelton DH80106-44N) that was forward biased through RF chokes (CoilCraft 1812CS-102XJEB, 1 μH, self-resonant frequency: 310 MHz) during transmission.

The Tx dipoles were installed and tuned with the Rx array already adjusted. The coil was placed inside a mock RF shield and loaded with the phantom to measure the Tx S-parameters with the Rx array detuned. This measurement was repeated with two volunteers. The S-parameters were later compared to the matrix obtained from the scanner RF safety watchdog log. Coupling to the Rx coil in the receive mode was also measured on the bench.

#### Phantom

For this study, we designed a 45-cm-tall head-neck phantom, with a 31-cm-wide section of the upper thorax (**Figure 1C**), by using the 3D average head model as a reference. The resulting head circumference was 60 cm, which is in the large range for an average adult male. Its inner volume was exported to CST to be used as the coil load during simulations. The phantom housing was 3D-printed in PLA (EcoTough™ PLA 2.0), waterproofed with an interior coating of epoxy (Smooth-On XTC-3D® High Performance Coating), and externally reinforced with the same epoxy and fiberglass (120-38 Standard E-Glass fiberglass cloth). It was filled with a solution based on published recipe^59^. The phantom contained the following ingredients per liter: demineralized water (398 ml), sugar (778 g), salt (106 g), alcohol-free mouthwash (40 ml), and Gadovist 1.0 (1 ml). A conductivity of 0.6 S/m and a relative permittivity of 35.5 were measured at 300 MHz with an Agilent 85070E dielectric probe. These parameters mimic the estimated average of the biological tissues that constitutes the region to which the coil is sensitive. This was verified on the bench by comparing reflection coefficients measured on different Rx loops loaded with the phantom and three volunteers. Finally, the phantom was used as a load during bench and scanner tests.

### Coil evaluation in the MRI room

#### Safety

The ethics committees of the scanning sites approved all in vivo 7 T MRI studies presented in this work. Examinations were performed on fifteen adult healthy volunteers who gave written informed consent.

MRI tests were performed in a MAGNETOM Terra scanner (Siemens Healthineers, Erlangen, Germany), using the parallel transmission (pTx) research mode. The integrity of coil components during high peak-power, high-SAR and high gradient-strength pulse sequences is critical to ensure the safe operation of the coil during routine use^60^. Coil heating was evaluated prior to and subsequent to a SAR-intensive sequence (operating at approximately twice the permissible SAR level for human scanning in first-level-controlled mode^61^) with an infrared camera. Commensurate temperature evaluation was conducted for high peak-power and gradient-intensive scans. No discernible increases in housing temperature were measured; therefore, the coil user is limited only by patient SAR (as described below) and not by the power-handling capacity of the coil.

#### Receive coil sensitivity profile

The sensitivity profile of individual receive coil elements was reconstructed from a sagittally oriented FLASH scan (in-plane FOV = 320 × 302 mm^2^, matrix size = 192 × 144, number of slices = 5, slice thickness = 5 mm, slice gap = 15 mm, TR/TE = 8/3.69 ms, FA = 20°, BW = 320 Hz/pixel).

#### Signal-to-noise ratio and geometry factor

SNR and geometry factor (g-factor) maps were acquired using a GRE scan. A sagittal scan with in-plane FOV = 384 × 384 mm^2^, matrix size = 256 × 256, number of slices = 13, slice thickness = 2 mm, slice gap = 0.4 mm, TR/TE = 30/6 ms, FA = 12°, averages = 2 and phase-encoding direction = superior-inferior, was acquired to map the g-factor along the superior-inferior and right-left directions. An axial scan was acquired to map the axial g-factor, with in-plane FOV = 192 × 192 mm^2^, matrix size = 384 × 384, number of slices = 7, slice thickness = 5 mm, slice gap = 5 mm, TR/TE = 30/6 ms, FA = 12°, averages = 2, and phase-encoding direction = right-left.

The scans were repeated without RF excitation to derive the noise covariance matrix. SNR maps were calculated using the root-sum-of-squares approach^9,62^, while g-factor maps were derived from the coil sensitivities and noise correlation information^21^. B_1_^+^ field effects on SNR were removed by dividing the SNR maps by the sine of the measured flip angle.

#### B_1_^+^ mapping and RF shimming

In vivo B_1_^+^ maps of individual transmit elements were acquired with a vendor-provided pre-saturation turbo-fast-low-angle-shot pulse sequence^63^ with the following parameters: FOV = 388 × 240 mm^2^, matrix size = 196 × 120 mm^2^, sagittal orientation, number of slices = 28, slice thickness = 3.2 mm, TR/TE = 6970/1.79 ms, saturation flip angle = 90°, excitation flip angle = 10°, BW = 550 Hz/pixel. These maps were combined offline using three different shim settings: (i) the nominal CP mode (i.e., a phase-only shim that produced the highest efficiency in C3); (ii) maximizing B_1_^+^ uniformity by minimizing the standard deviation over the mean B_1_^+^ along the spinal cord between the upper thorax and the occipital region; and (iii) maximizing the B_1_^+^ efficiency over the same region. Subsequently, the SAR efficiency of the B_1_^+^ field was calculated using the maximum 10-g-averaged SAR derived from the local SAR matrices (excluding the 2-fold safety factor utilized for human scanning). All B_1_^+^ shimming calculations were performed in MATLAB.

An MP2RAGE^64^ scan was acquired in CP mode and sagittal orientation, FOV = 256 × 224 × 192 mm^3^, 1 mm^3^ isotropic resolution, TR/TE = 3250/1.83 ms, TI1/TI2 = 840/2370 ms, FA1/FA2 = 5/6°, in-plane acceleration factor = 3, BW = 250 Hz/pixel.

The spinal cord between C1 and T5 was segmented from the GRE image of the second inversion time (INV2) of the MP2RAGE scan using *Spinal Cord Toolbox*^*65*^ and used as a mask for B_1_^+^ shimming as described below.

Using *Shimming Toolbox*^66^, a set of eight complex shim weights (***w***) was calculated offline, such that the resulting B_1_^+^ efficiency would be uniform across the spinal cord:

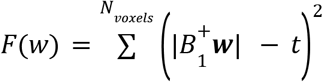

where F(***w***) is the cost function to be minimized via B_1_^+^ shimming, *t* is the target B_1_^+^ efficiency of 15 nT/V, B_1_^+^ is the B_1_^+^ maps for all Tx channels and *B*_1_^+^***w*** is the B_1_^+^ map for all Tx channels after the shim weights, ***w***, are applied. Shim-weight optimization was performed using the *Shimming Toolbox*. During the optimization process, local SAR was constrained and set to not exceed the maximum local SAR obtained with the CP mode by more than 50%—local SAR was computed using VOPs^57^.

Using the shim weights, ***w***, B_1_^+^ maps and a new MP2RAGE scan were re-acquired and masked using the spinal cord mask described above. Signal intensity, T1 value and B_1_^+^ efficiency within this mask were analyzed for the MP2RAGE INV2 images, MP2RAGE T1 maps and in-vivo B_1_ ^+^ maps, respectively, along with the coefficient of variation (CoV).

#### Comparison with other coils

An identical coil was built and supplied to a second site (MGH Martinos Center for Biomedical Imaging, USA) to perform multicenter studies. This coil version was compared to two different coils^67^: an 8-Tx/32-Rx brain coil (Nova Medical, Wilmington, MA, USA) and a custom-built 16-Tx/64-Rx head-neck coil^5^. SNR quantitative maps were calculated by means of a root-sum-of-squares reconstruction with noise-covariance weighting of the individual channel data^9,62^. A gradient-echo sequence was used (TR = 8000 ms, TE = 3.82 ms, FOV = 256 mm, voxel size = 1 × 1 × 2 mm^3^, number of slices = 130, FA = 78°, bandwidth = 340 Hz/px, reference amplitude = 500 V and 0 V). For each coil, the optimal SNR combination was then divided by the sine of the flip angle to remove variations due to transmit-field non-uniformities^68^. A region-of-interest was manually-drawn on the acquired images to calculate the SNR profiles along the z-axis from the spine to the brain.

## Results

### Receive coil

With the Rx coil completely adjusted and loaded with the phantom, the mean output impedance measured at the input sockets of the preamplifiers, towards the loops, was 75.3 ± 3.9 Ω; the reactance, which was reduced to a minimum during the adjustments showed a mean of -0.6 ± 0.3 Ω. A confidence level of 95% was used here and in all the following data. Coupling (S_nm_) measured under the same loading conditions showed a close agreement with values predicted by simulation. **Table 1** shows the mean coupling values between adjacent loops and the overall coupling obtained separately from each subarray. The simulated average coupling between both subarrays was -32.4 ± 2.4 dB; only simulations were used in this case because measurements on the independent sections showed that the CST model of the coil was accurate. As expected, the highest coupling (≈ -7 dB) was obtained, and later verified by measurements, from pairs of loops 8-17 and 12-16. These are the elements from different subarrays that are positioned at the closest distance, without being critically overlapped (see **Figure 2C**). The mean value of measured preamplifier decoupling was -13.8 ± 2.6 dB.

**Table 1.** Mean coupling values showing good agreement between bench measurements and initial simulations. In both cases the phantom was used to load the coil. Adjacent coupling of -15 dB is considered satisfactory when preamplifier decoupling is implemented.

| Mean coupling, $S_{nm}$, in dB | | | | |
| --- | --- | --- | --- | --- |
|  | 15-Ch (posterior) |  | 5-Ch (anterior) |  |
|  | *Measured* | *Simulated* | *Measured* | *Simulated* |
| Adjacent | $-15.6 \pm 0.9$ | $-15.8 \pm 2.4$ | $-16.3 \pm 1.3$ | $-17.2 \pm 1.9$ |
| Overall | $-22.3 \pm 1.7$ | $-23.6 \pm 2.1$ | $-16.6 \pm 1.2$ | $-17.3 \pm 1.4$ |

Images acquired with the individual channels, shown in **Figure 4**, demonstrated the coverage over the region of interest and limited coupling between Rx elements. **Figure 5** shows in vivo SNR maps over the central sagittal plane and an axial plane located at the C3-C4 segment. The SNR along the spinal cord increases from the thoracic to the occipital region, which is in agreement with the increase in density and proximity of receive elements. **Figure 6** shows the inverse g-factor (1/g) maps where typical values for 1D and 2D accelerations were obtained from slices that will be regularly selected during spinal cord studies. Noise enhancement in the spinal cord is in an acceptable range up to a one-dimensional acceleration factor of 4. As for two-dimensional acceleration, less desirable noise amplification values start to appear from R = 3 × 3.

**Figure 4.**
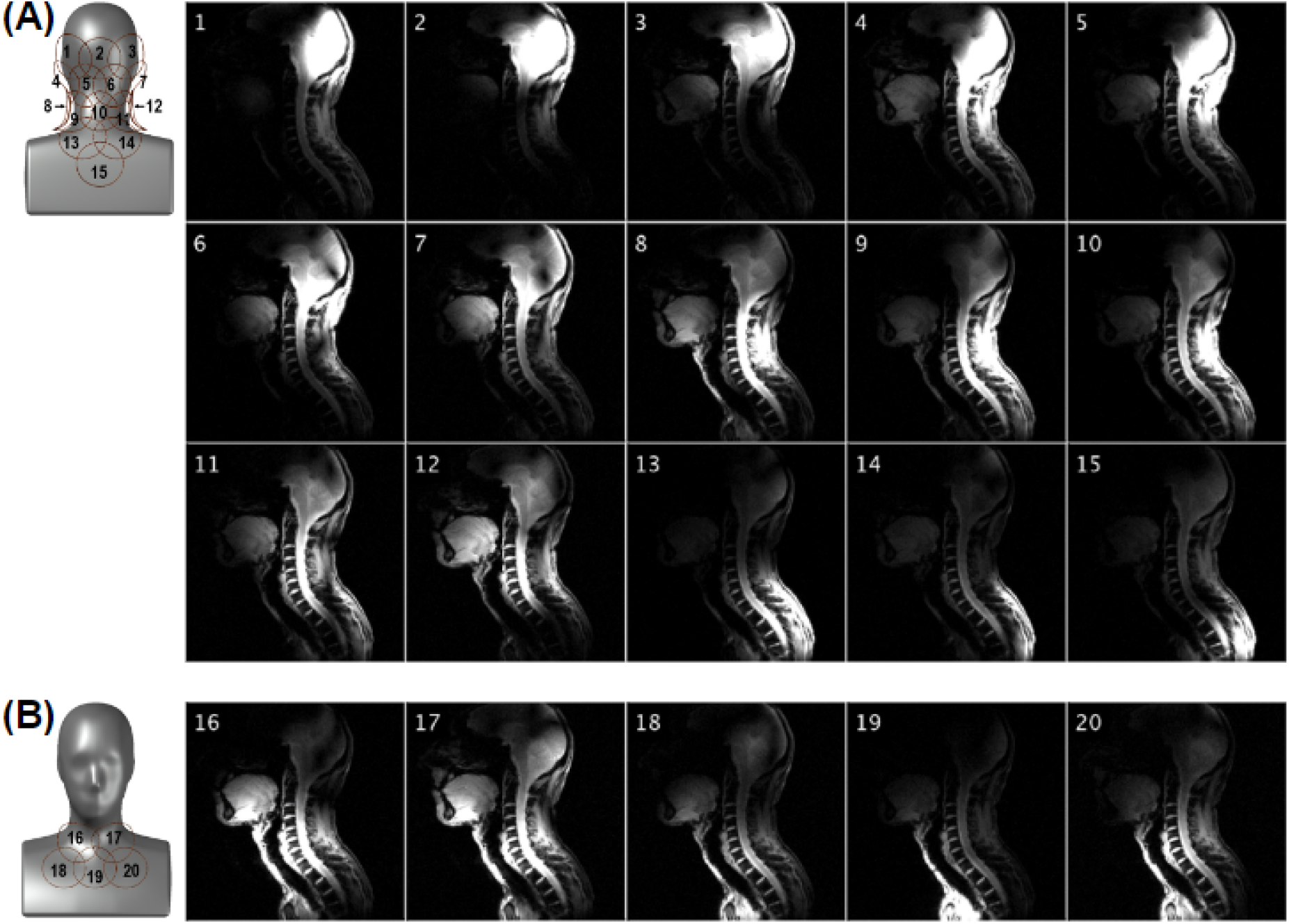
Sensitivity profile of the posterior **(A)** and anterior **(B)** Rx subarrays obtained with a FLASH sequence having an FOV of 320 × 320 mm and a 512 × 512 matrix. The intensity colormap scaling was kept the same across the 20 panels.

**Figure 5.**
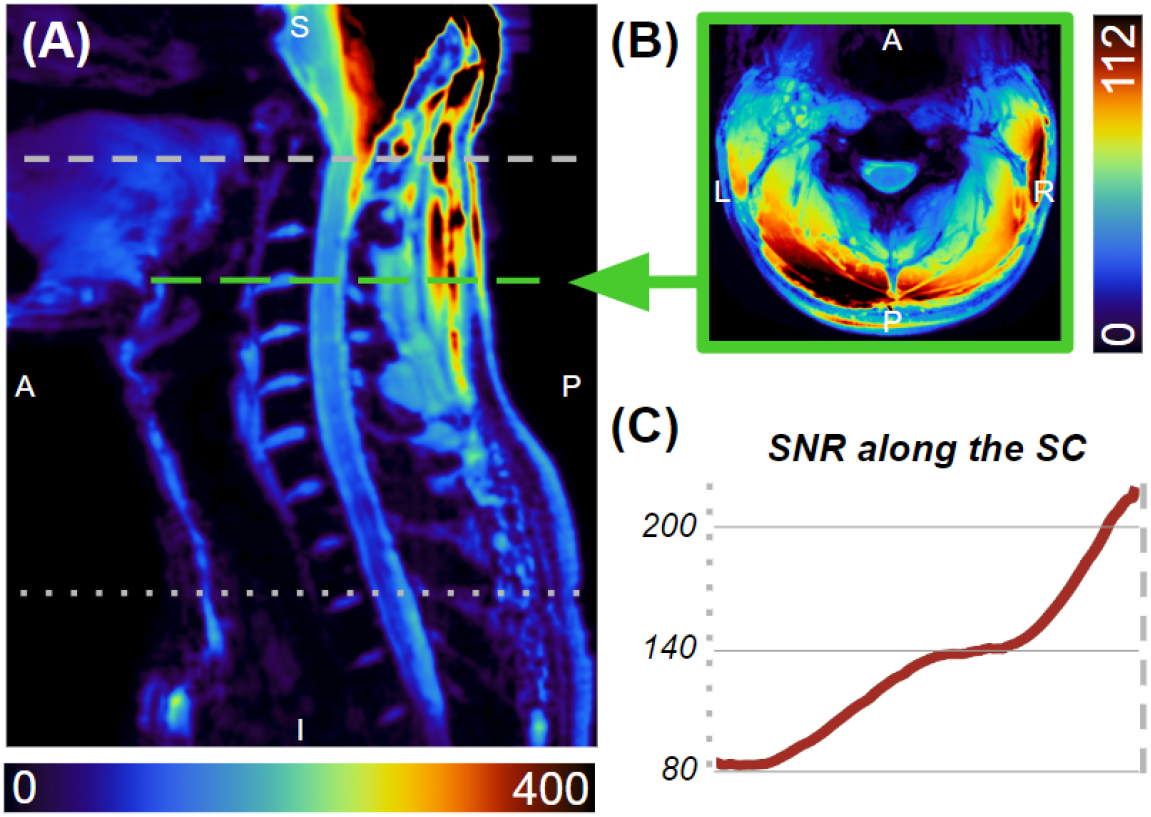
SNR maps along the sagittal midline **(A)** and the C3-C4 level **(B)**. SNR along the spinal cord **(C)** ranges from 80 at the T2-T3 disc, increasing to 200 at the top of the C1 vertebra.

**Figure 6.**
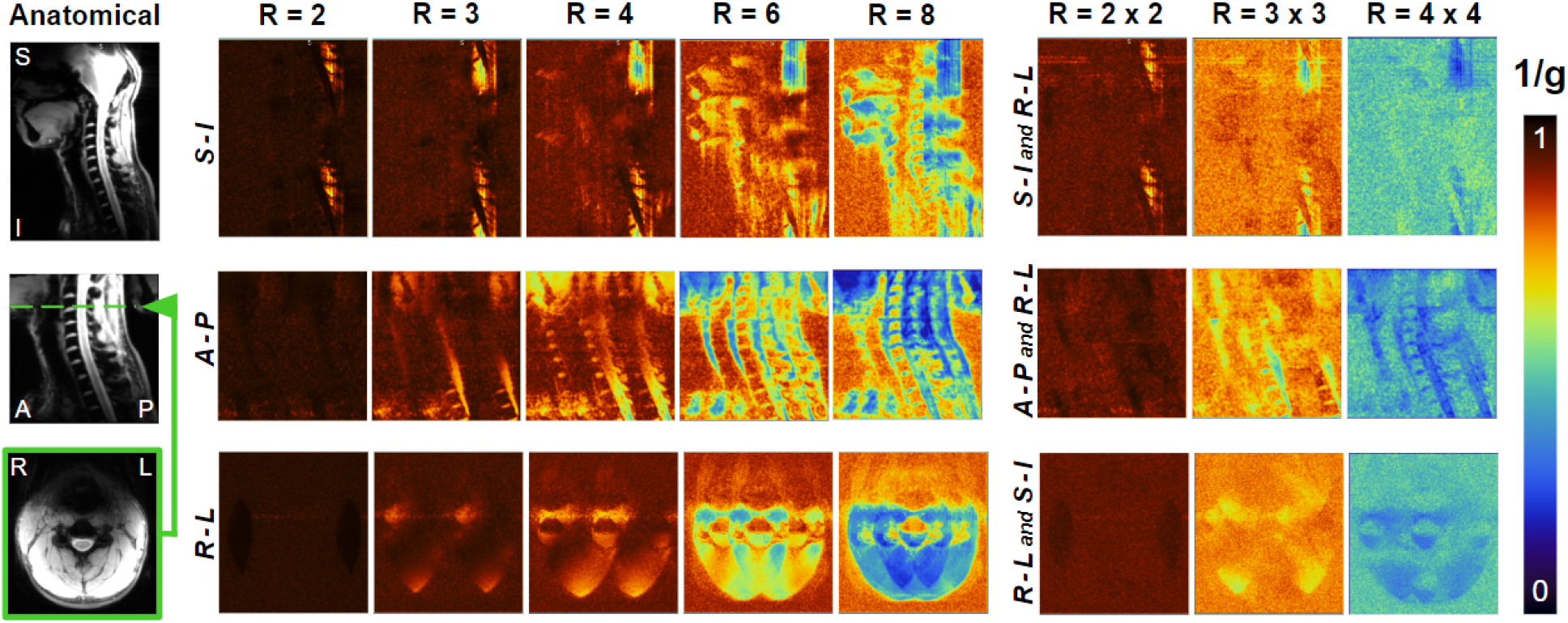
Maps of the inverse g-factor (1/g), show at the sagittal midline with acceleration in the superior-inferior and right-left directions (top), at the sagittal midline with acceleration in the anterior-posterior and right-left directions (middle), and axially at the C3-C4 level with acceleration factors in the right-left and superior-inferior directions (bottom).

### Transmit coil

The mean impedance of bazooka baluns and RF chokes was 710 Ω and 1130 Ω, respectively. No sensitivity to touching or repositioning of the transmit cable was observed, which demonstrated that the combination of a bazooka balun and RF choke was sufficient to achieve common mode suppression.

The maximum and mean coupling between adjacent transmit elements, when measured on the bench, inside the mock RF shield, with the coil loaded by the phantom and the Rx-coil detuned, were -7.7 dB and -16.5 ± 6.5 dB, respectively. The mean coupling between all elements was -23.4 ± 2.8 dB. The mean reflection was -32.6 ± 2.3 dB. The corresponding S-parameter matrix is shown in **Figure 7A**. A similar matrix was obtained from the scanner with the same phantom (**Figure 7B**). In this case, the maximum and mean coupling between adjacent transmit elements were -7.9 dB and -15.6 ± 6.3 dB, respectively. The mean coupling between all elements was -23.8 ± 3.5 dB. The mean reflection was -23.5 ± 2.3 dB. The adequate similarity observed between the two matrices validated the adjustments performed on the bench. Active detuning provided a mean isolation of -21.5 dB ± 2.3 dB with the receive coil during reception.

**Figure 7.**
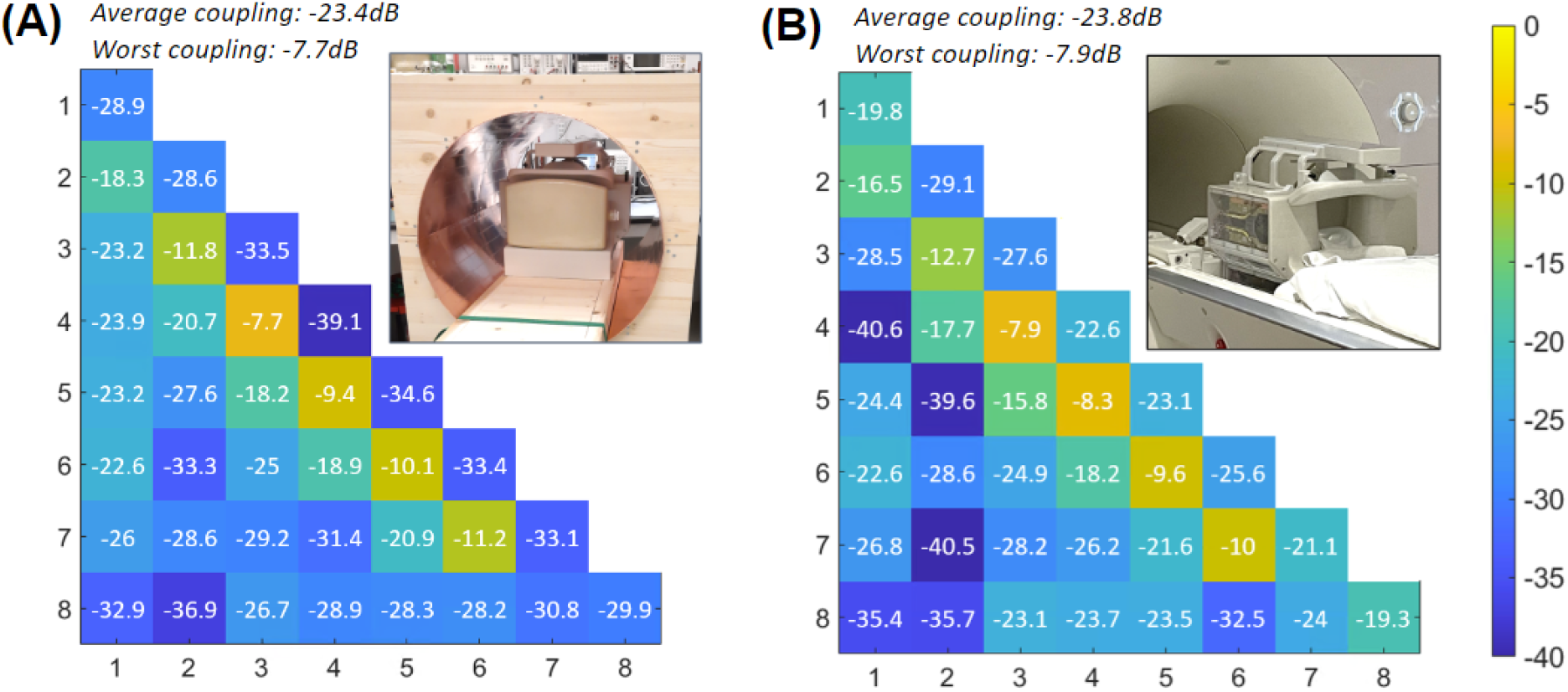
S-parameter matrices of the Tx coil measured on the bench inside a mock RF shield (**A**) and obtained from the scanner (**B**), showing a fair resemblance.

Simulated B_1_^+^ efficiency and 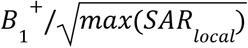 is shown in **Figure 8** for the four CST body models incorporated into the VOPs. The maximum 10-g-averaged SAR is located in the nose for a large human model, like Hugo, due to the close proximity of the anterior dipole. For small-to-average size human models, the maximum local SAR occurs in the neck region.

**Figure 8.**
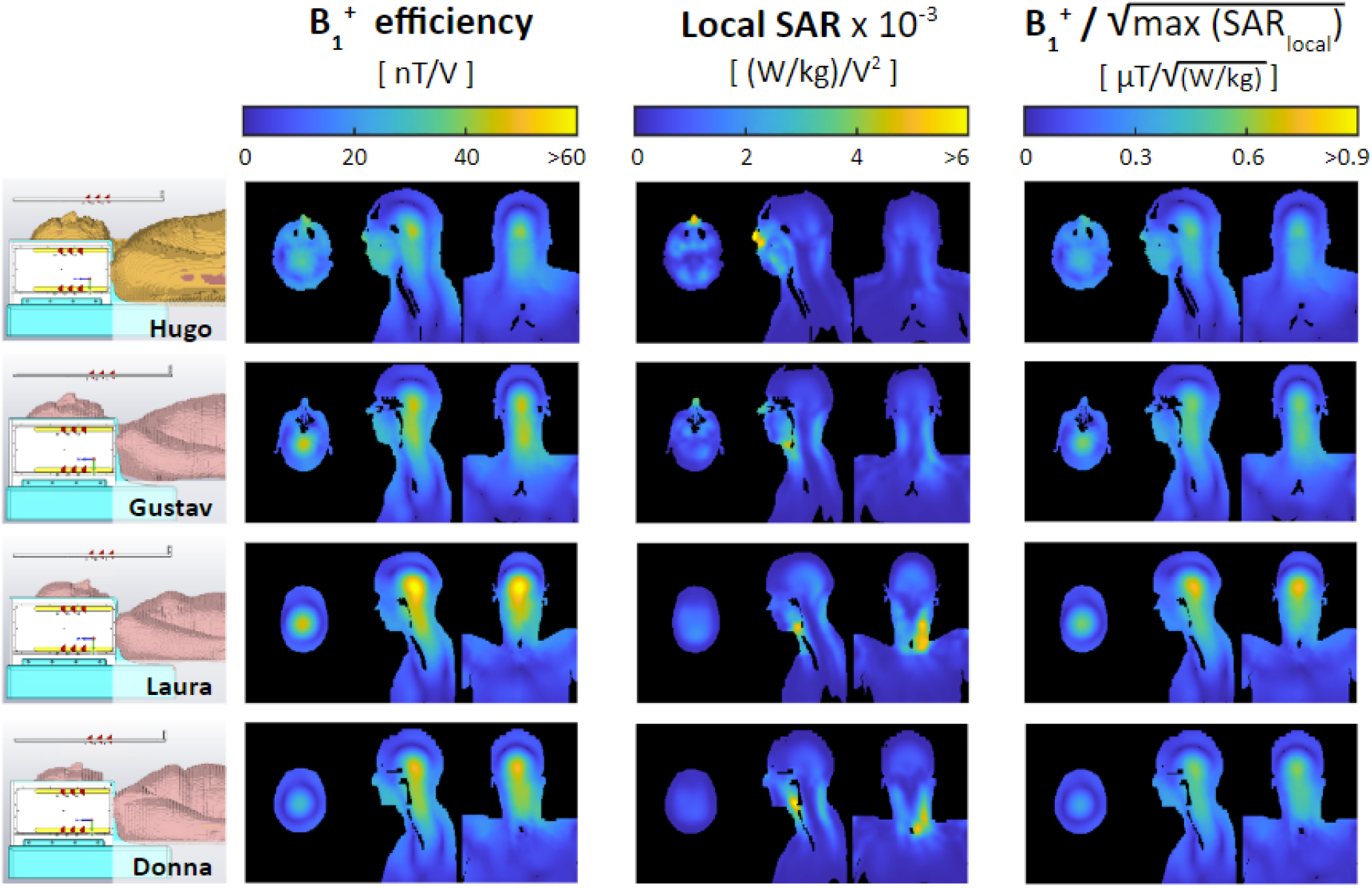
Simulated B_1_^+^ efficiency, 10-g-averaged SAR efficiency, and B_1_^+^ efficiency per square root of the maximum local SAR for four CST body models (shown on the left)—Hugo, Gustav, Laura, and Donna—when driven in the nominal CP mode. The maximum local SAR occurs in the nose for a large body model like Hugo (due to close proximity to the anterior transmit dipole) and in the neck for small-to-average-sized body models. All four body models were incorporated into online SAR matrices—the diversity of which ensures a conservative estimate of local SAR.

### B_1_^+^ shimming

RF shimming capabilities of the coil were demonstrated by simulations performed on the validated coil model (see Supporting information). Implementation examples are shown in **Figure 9**. In all cases, the spinal cord was segmented using the *Spinal Cord Toolbox* and overlapped on the original images. A comparison of two B_1_^+^ efficiency combinations generated from individually acquired B_1_^+^ maps is shown in **(A)**: in CP mode (left) and with optimized shim weights (center). **Figure 9 (B)** shows the signal intensity of GRE images from the second inversion time, while **Figure 9 (C)** shows the estimated T1 values. The shim weights were optimized to minimize the CoV along the spinal cord.

**Figure 9.**
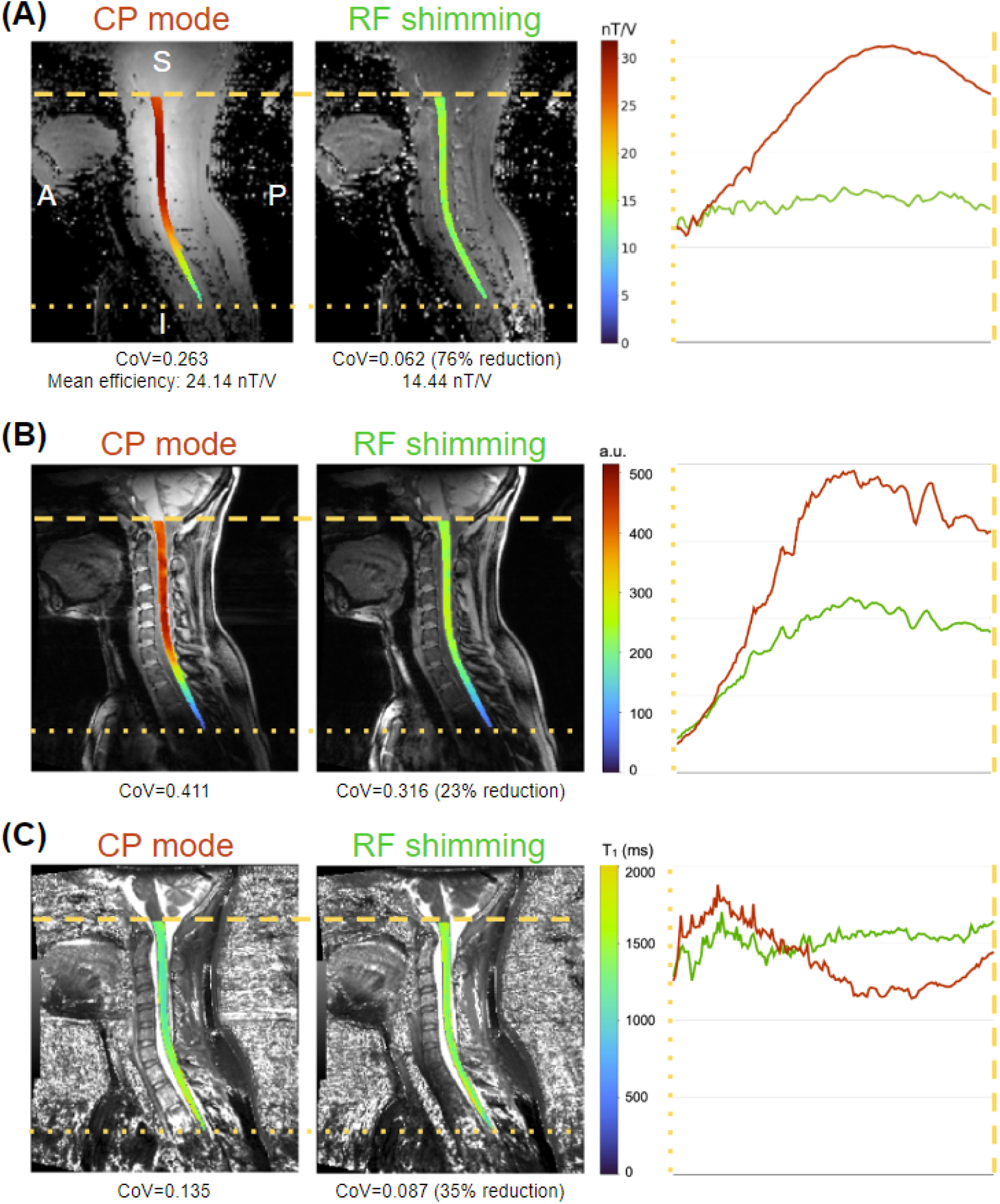
Example of the effect of RF shimming for optimal uniformity on the B_1_^+^ map (A), GRE scan (B) and T1 maps from the MP2RAGE sequence (C).

CP mode excitation resulted in an inhomogeneous excitation profile, with high B_1_^+^ efficiency and corresponding high signal intensity at the upper cervical levels, with a steep drop from ∽C4 to ∽T2, as shown in the left panels of **Figure 9 (A)** and **(B)**. RF shimming resulted in a more homogeneous excitation profile, evidenced by a reduction of CoV by ∽76% (from 0.263 to 0.062) for B_1_^+^ efficiency and 23% (0.411 to 0.316) for GRE signal intensity. The measured B_1_^+^ efficiency for RF shimming (14.44 nT/V) is in line with the targeted B_1_^+^ efficiency of 15 nT/V.

Improved B_1_^+^ homogeneity resulted in more uniformity along the spinal cord for estimated T1 values, evidenced by a CoV reduction of ∽35% (0.135 to 0.087). Additionally, while the T1 values estimated in the CP mode show a marked reduction in the upper cervical levels (and a similar increase in the upper thoracic spine), T1 values estimated from the RF shimmed MP2RAGE scan are relatively uniform along the FOV.

### Comparison with other coils

**Figure 10** shows an SNR comparison of the proposed coil and the two available alternatives^67^. The SNR map revealed adequate uniformity (CoV = 10.2%) from the occipital lobe to the mid-thoracic spine, while showing a 30.4% increase in mean SNR (SNR_head/neck_ = 246.9, SNR_head_ = 90.9, SNR_spine_ = 321.9) with respect to the head-neck, which was designed to include the spine region as well. The SNR declines in the upper part of the studied region, which comprises most of the brain, but useful values were registered in the occipital region.

**Figure 10.**
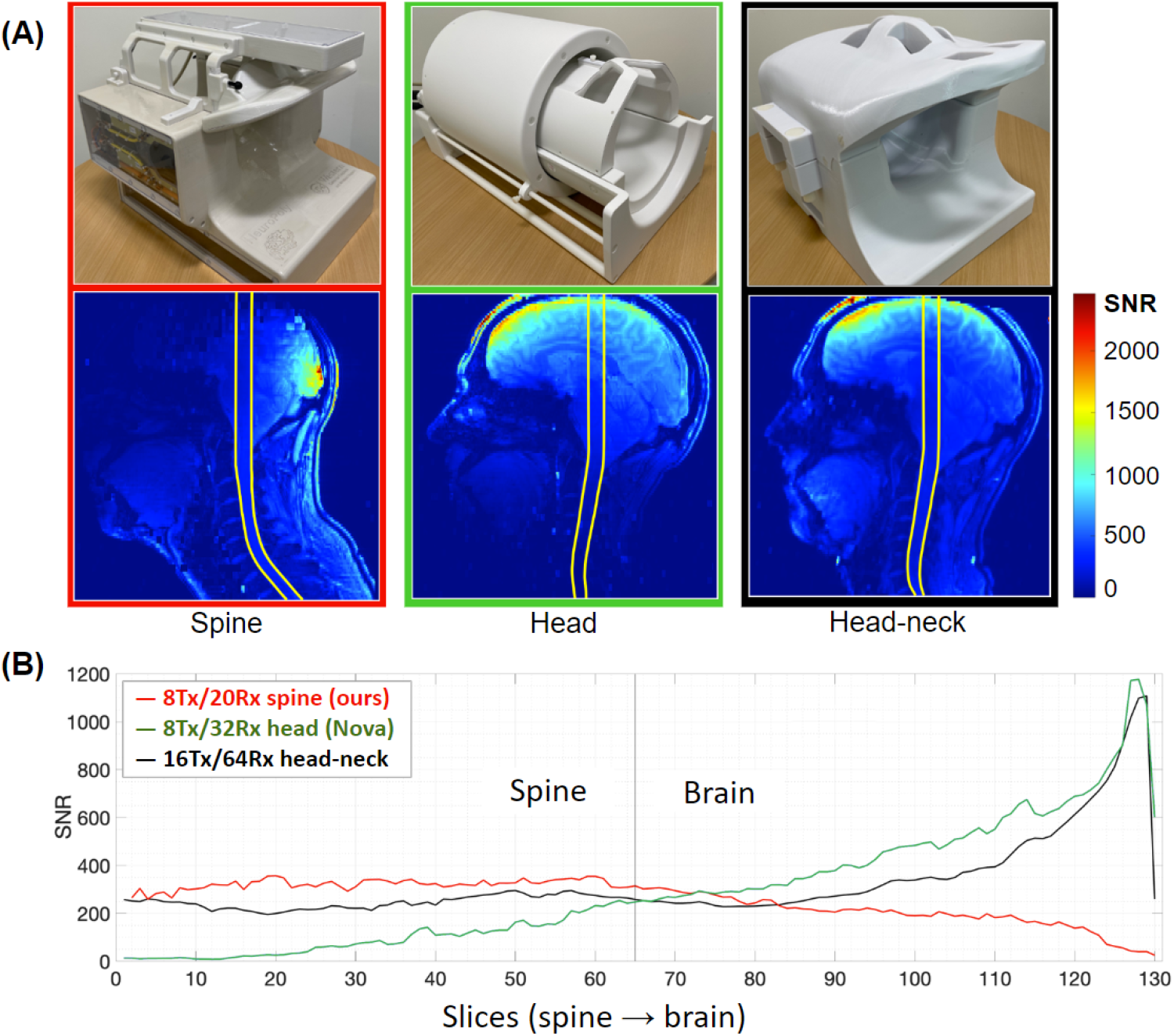
SNR assessment and comparison with two different coils displaying SNR maps **(A)** and profiles **(B)**. Each point on the profiles is the average SNR within the region of interest shown on the corresponding maps for each slice^67^.

## Discussion

The development of MRI coils to study a region extending from the occipital lobe to the T4-T5 vertebral level at 7 T has to overcome several challenges to generate a uniform excitation field and achieve high sensitivity. Most of these challenges are caused by the length, curvature and location of the spinal cord itself, and the complex anatomy of this region—characterized by a variety of tissues, dimensions and morphological differences within the population. In this work, we present an 8-channel transmit and 20-channel receive coil that addresses these challenges. Testing of this coil showed excellent performance over an extended range beyond the cervical spine including the occipital and upper thoracic regions when compared to other coils^5,6^. The Tx and Rx coils are integrated into a single housing equipped with hinges that allow it to be easily opened and closed to expedite the placement of the subject. In addition, the position of the anterior portion of the receive coil can be adjusted to better conform to the anatomy of the subject.

### Rx coil

The difficulties involved in conceiving a receive array adapted to this complex anatomical region and the need to know the exact geometry of the receive coil for future experiments motivated the use of CST simulations to carry out the design task. The time spent on the simulations was compensated by the simplification of the assembly and adjustment, especially of the critical overlap between adjacent elements, which is also a time-consuming task. In this regard, it was not necessary to empirically reshape the wire loops once they were mounted directly into the grooves created on the coil formers, since the inductive decoupling values achieved in the simulations were preserved.

The mean coupling (−16 dB) was also close to the recommended level when preamplifier decoupling is implemented^69^. In some cases (loops 5-6, 9-10, 6-12, 16-17), the best possible S_nm_ values were between -10.6 and -12 dB, mainly due to loop shapes that favored coupling. The worst overall values (−7.1 dB), obtained for loop pairs 8-17 and 12-16, were caused by the proximity of the respective elements when the anterior (adjustable) array section was placed at its closest position to the fixed posterior to better adapt the array to the phantom; however, these values improved when the array was adjusted for a larger load, since elements 16 and 17 are moved away from 12 and 8, respectively. In all cases, the proximity between the simulated and measured coupling values confirmed the validity of the simulations.

Comparison to other coils (head and head-neck) with more channels demonstrated that 20 receive loops, with convenient dimensions and placement, were sufficient to obtain higher SNR along the spinal cord^5^. Average g-factors obtained during the same study over an axial plane positioned at level C3 of the cervical spine showed similar values for all coils up to acceleration factors (R) of 4, while reductions of 27% (0.67/0.49) and 22% (0.63/0.49), respectively, were observed for R=6. This is an expected result mainly due to the lower number of Rx channels of the proposed coil. However, even though one of the initial objectives was to build a coil with a minimum number of channels, both aspects could benefit slightly from the addition of a Rx element near the upper thorax without significantly increasing the complexity of the coil.

### Tx coil

The cervical spinal cord is located considerably deep into the body and has a characteristic curvature that causes a variable distance to the elements of any coil along the Z-axis. This combination presents a difficulty in attaining high power efficiency and entails the use of RF shimming to produce a homogeneous excitation field at 7 T. The presence of the shoulders adds disparate geometrical constraints along Z. They prevent the use of a volume coil that surrounds the spinal cord closely enough to operate with acceptable efficiency and with sufficient length to cover our desired region of interest (down to the upper thorax).

Dipoles were selected for the Tx coil primarily for the ability of their geometrical properties to address the specific challenges described above. The physical length of loaded, half-wavelength dipoles at 7 T (approximately 44 cm when near the head^46^) allowed for a transmit coil to be developed with an extended B_1_^+^ coverage along the superior-inferior axis, despite being limited to only eight transmit channels (a ubiquitous specification for ultra-high field scanners at present). The holistic approach in designing the receive array resulted in high decoupling between both coils. This was highly influenced by the selection of non-uniform spacing between transmit dipoles, which created sufficient clearance to place the preamplifiers and other receive components. As a result, the tuning process of both coils was significantly simplified, as it was possible to perform them independently. In this way, we avoided the difficulties inherent to Tx/Rx coils with highly-coupled and closely-spaced arrays, as encountered in the early stages of other works^3,5^, while protecting the Rx coil during the Tx mode and potentially reducing local SAR.

With each individual element being sensitive to a larger region of the body, a volume-like multi-channel topology could be implemented, while still maintaining close proximity to the region-of-interest. The resultant higher field intensities in the central region (i.e., central brightening), characteristic of volume coils, increased the efficiency around the spine. Volume coils, in comparison to planar coil arrays, are compatible with automatic and convergent transmit-power calibration when scanning—a significant advantage for clinical workflows. The multi-channel design could then be used for RF shimming to further improve the the performance of the coil over an extended field-of-view.

RF shimming over the longitudinally oriented spinal cord was further benefited by the difference and offset of the B_1_^+^ profiles of the lateral dipoles with respect to those located in the posterior and anterior sections of the coil. As a consequence, the FOV along the superior-inferior axis was lengthened, which made it possible to study an elongated ROI extending from the occipital lobe to T4-T5 vertebral levels. The results obtained in this region showed higher power efficiency and lower SAR than other solutions where pTx was not implemented^6^. In addition, RF shimming can provide a range of solutions depending upon the application. Simulations showed that the coil offered the flexibility required to implement B_1_^+^ shimming solutions in this region to improve transmission uniformity, efficiency, or SAR according to the requirements of the pulse sequence. Trade-offs between these parameters can also be obtained by modifying the regularization of the cost functions.

In conclusion, this work presents a transmit/receive coil, composed of 8 pTx dipoles and 20 anatomically shaped semi-adaptable Rx loops for spinal cord imaging at 7 T. The coil was designed, tested and validated by simulations, on the bench and in the scanner. RF shimming allowed optimization of B_1_^+^ uniformity, B_1_^+^ efficiency or SAR efficiency. An extended field of view covering from the occipital lobe to the upper thoracic spine was demonstrated with several volunteers. A comparison with available coils revealed improved performance parameters in the spinal cord region. This coil solution therefore provides a convincing platform for producing the high image quality necessary for clinical and research scanning of the upper spinal cord.

## Acknowledgments

The authors thank Justin De Meulemeester for helping with VOP creation, Dr. Pedram Yazdanbakhsh and the 7T team at the Brain Imaging Center of the Montreal Neurological Institute for technical support during imaging, Dr. Marcus Couch, Dr. Omer Oran and Dr. Yulin Chang from Siemens Medical Solutions for technical support. This study was funded by the Canada Research Chair in Quantitative Magnetic Resonance Imaging [950-230815], the Canadian Institute of Health Research [CIHR FDN-143263], the Canada Foundation for Innovation [32454, 34824], the Fonds de Recherche du Québec - Santé [322736], the Natural Sciences and Engineering Research Council of Canada [RGPIN-2019-07244], the Canada First Research Excellence Fund (IVADO and TransMedTech), the Quebec BioImaging Network [5886, 35450] and Mila - Tech Transfer Funding Program. K.M.G. received financial support from the Canada Foundation for Innovation, Canada First Research Excellence Fund to BrainsCAN, and Brain Canada Platform Support Grant. Imaging performed at the Athinoula A. Martinos Center for Biomedical Imaging used resources provided by the Center for Functional Neuroimaging Technologies (P41EB015896) and the Center for Mesoscale Mapping (P41EB030006), Biotechnology Resource Grants supported by the National Institute of Biomedical Imaging and Bioengineering, National Institutes of Health (NIH). The NIH also provided support through a shared instrumentation grant S10OD023637 (L.L.W.) and grant R01EB027779 (R.L.B.). The content is solely the responsibility of the authors and does not necessarily represent the official views of the NIH.

## Data availability statement

The CAD design, PCB of dipoles, socket board for preamps, CST project for EM simulation are available at: https://github.com/neuropoly/coil-spine7t

## Supporting information

### Simulated B_1_^+^ fields

The simulated coupling between adjacent transmit elements (when converted to a fraction of coupled power) matched experimental measurements to within 2 percentage points: this close agreement ensured consistency between the spatial patterns of the corresponding B_1_^+^ distributions (**Figure 11**). The mean and 95^th^ percentile simulated and experimental B_1_^+^ maps of the phantom, when combined in CP mode, differed by -11.5% and 1.4%, respectively. For 10,000 random shim settings (magnitude and phase), the vast majority of solutions resulted in deviations between any combined simulated and experimental B1+ map that ranged within +/-40% (mean and 95th percentile). However, these results included some uncertainty because the masking of the simulated and experimental maps were different.

**Figure 11.**
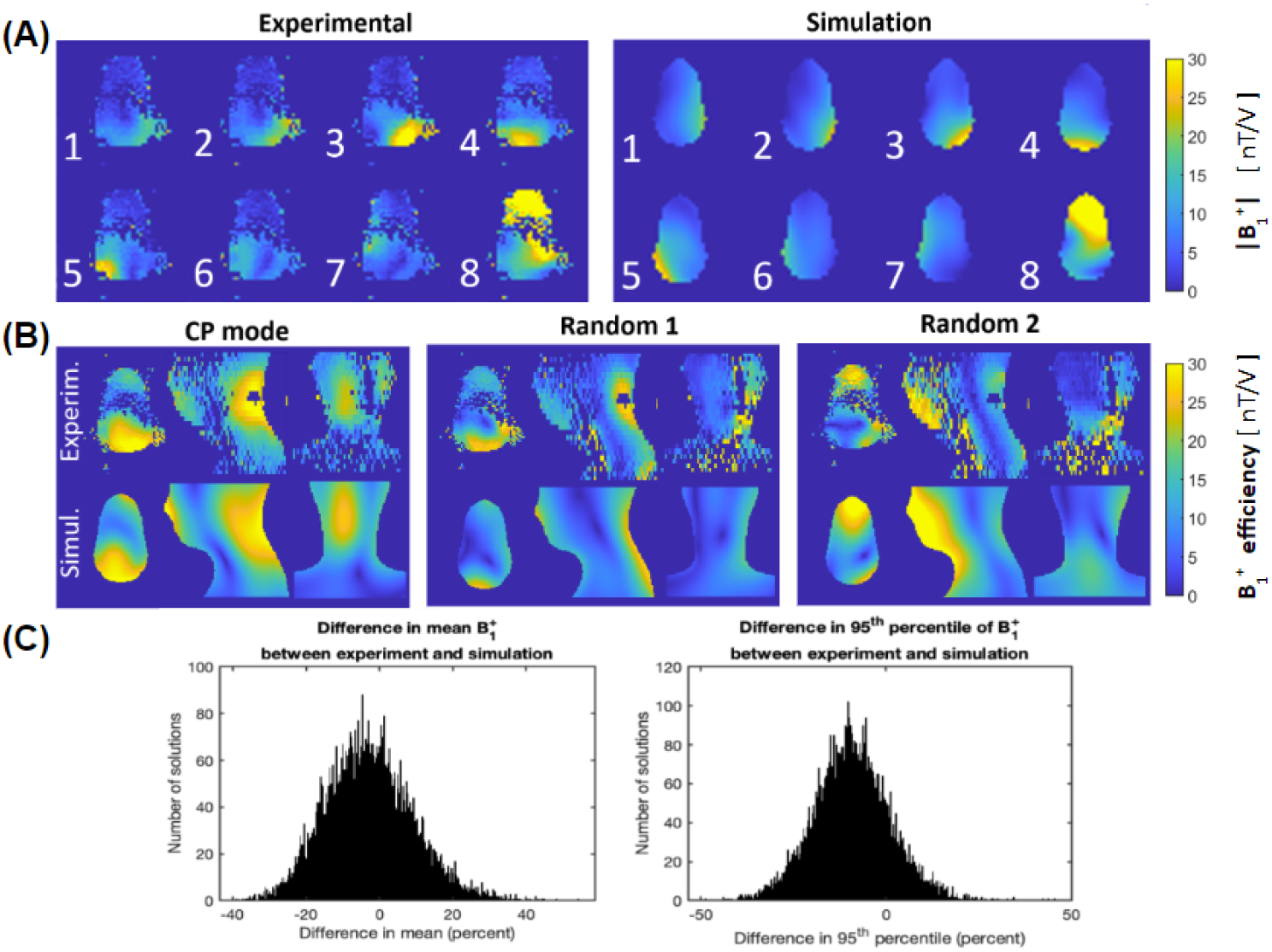
**(A)** A representative axial slice of B_1_^+^ maps (of the spine phantom) from individual transmitters derived from experiment (left) and simulation (right). **(B)** Complex scaling factors were applied to simulated B_1_^+^ fields to increase agreement with experimentally derived maps—resulting in a commensurate agreement when comparing B_1_^+^ maps combined in CP mode (left) and two random shim modes (center, right). **(C)** When combining the eight transmit channels with 10,000 different random shim settings, histograms of the deviation between the mean and 95th percentiles of the simulated and experimental B_1_^+^ distributions showed that most of the solutions are within +/-40%.

### Simulated B_1_^+^ shimming

B_1_^+^ shimming provides the flexibility to tailor the B_1_^+^ field according to the requirements of the pulse sequence, whether that necessitates an improvement in transmit uniformity, efficiency, or local SAR. Individual B_1_^+^ maps, when combined in the nominal CP mode (phase-only shim solution), produce a mean B_1_^+^ over the spinal cord of 17.9 nT/V with a CoV of 39.8% (see **Figure 12 (A)**). When a shimming algorithm that maximizes transmit uniformity is employed, this metric is improved to 15.4% with a reduction in mean B_1_^+^ of 69%. If the transmit efficiency is maximized, a phase and magnitude shim solution can improve the efficiency of the CP mode by 12%, with a 2.8% decrease in uniformity. Shimming solutions can be computed within the bounds of these solutions by altering the regularization of the cost function, which allows a trade-off between B_1_^+^ efficiency, uniformity, and local SAR.

**Figure 12.**
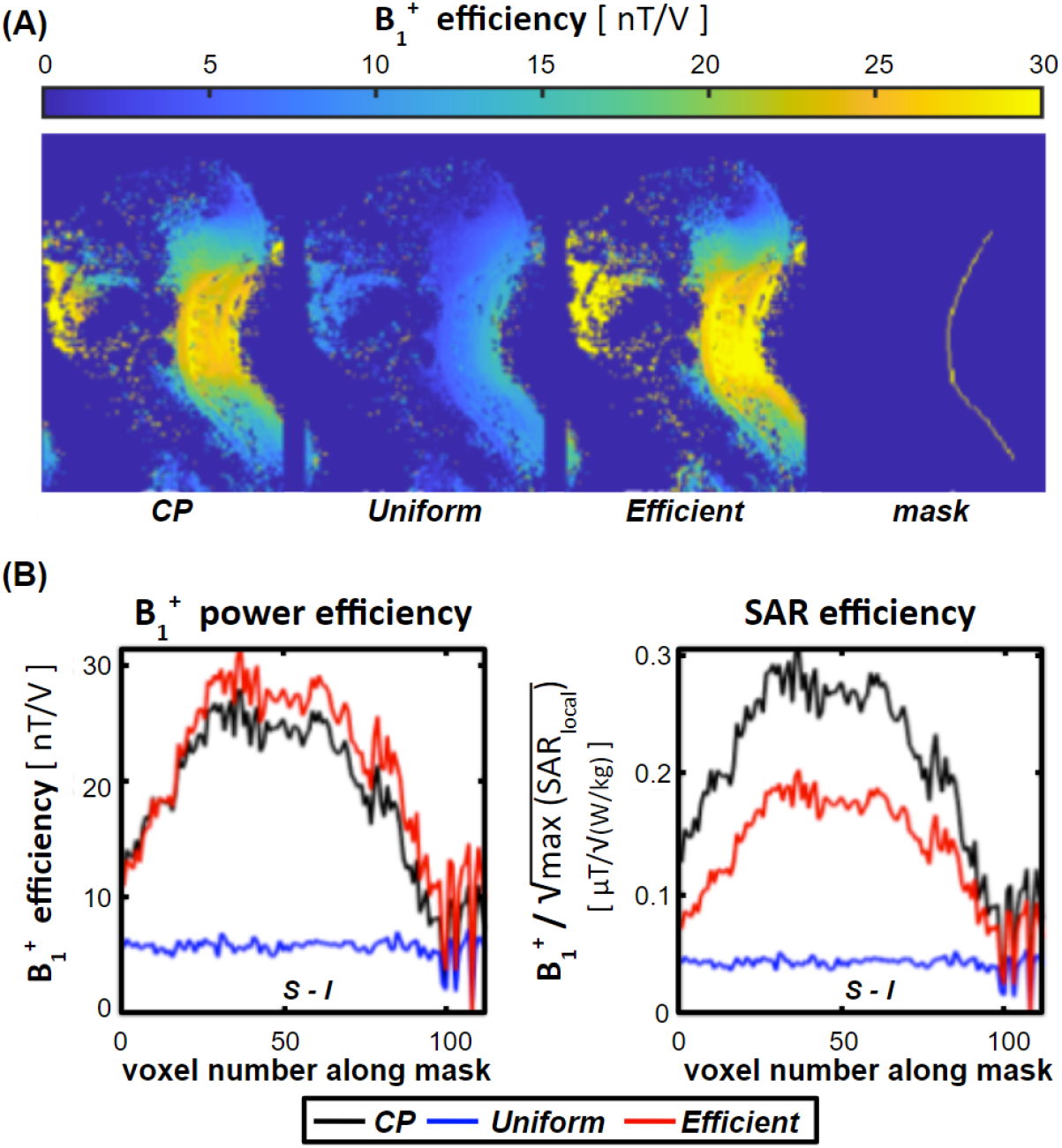
**(A)** B_1_^+^ maps with transmitter shim weights set to produce: (left) the nominal CP mode (i.e., a phase-only shim that produces the highest B_1_^+^ efficiency in the C3 vertebrae); (center) the highest B_1_^+^ uniformity over the spinal-cord region defined by the mask; and (right) the highest B_1_^+^ efficiency over the same region. **(B)** Plots of the B_1_^+^ power efficiency and SAR efficiency for these shim solutions demonstrate the breadth of the solution space—the shimming algorithm can then be tailored to the requirements of the desired pulse.

Whereas high transmit efficiency may be required for short, high flip-angle RF pulse, the SAR efficiency of the shim solution 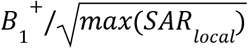, is often the limitation when determining the allowable number of slices and repetition time. Power efficiency and SAR efficiency are not necessarily synonymous, as evidenced by the 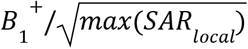 of the shim solutions presented in **Figure 12 (B)**, where the CP mode attained the highest 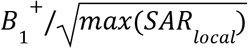, but not the highest power efficiency.

